# PoPoolationTE2: comparative population genomics of transposable elements using Pool-Seq

**DOI:** 10.1101/038745

**Authors:** Robert Kofler, Daniel Gómez-Sánchez, Christian Schlöetterer

## Abstract

The evolutionary dynamics of transposable elements (TEs) are still poorly understood. One reason is that TE abundance needs to be studied at the population level, and despite recent advances in sequencing technologies, characterizing TE abundance in multiple populations by sequencing individuals separately is still too expensive. While sequencing pools of individuals (Pool-Seq) dramatically reduces sequencing costs, a comparison of TE abundance between pooled samples has been difficult, if not impossible, due to various biases. Here, we introduce a novel bioinformatic tool, PoPoolationTE2, which is specifically tailored for the comparison of TE abundance among pooled population samples or different tissues. Using computer simulations we demonstrate that PoPoolationTE2 not only faithfully recovers TE insertion frequencies and positions but, by homogenizing the power to identify TEs acrosss samples, it provides an unbiased comparison of TE abundance between pooled population samples. We anticipate that PoPoolationTE2 will greatly facilitate the analysis of TE insertion patterns in a broad range of applications.

## Introduction

Transposable elements (TEs) are short stretches of DNA that selfishly propagate within genomes and are thought to be involved in diverse phenomena ranging from human diseases (Kazazian, 1998) to genome evolution (Kazazian, 2004).

Addressing many questions about the biology of TEs requires comparing TE abundance between different samples. Promising lines of research include the activity of TEs in mutation accumulation lines (e.g. base population vs mutated lines), the dynamics of TE invasions during experimental evolution (e.g. evolved populations at different time points), the contribution of TEs to local adaptation (e.g. populations from different areas), the evolution of TE activity (e.g. populations from different species) and the extend of somatic TE activity (e.g. different tissues) (González et al., 2008; Kofler et al., 2015; Perrat et al., 2013). Sequencing individuals (cells) separately may for many of these questions either be too costly or, as in the case of tissues, technically too challenging (sequencing of single cells), but sequencing pools of individual offers a viable alternative approach [Pool-Seq Schlötterer et al. (2014)]. However, a comparison of TE abundance between pooled samples is difficult as numbers of reads are usually insufficient to identify all TEs within a pool. This leads to a clear bias, with most TEs being frequently found in the sample with the most mapped reads. While it is possible to standardize the number of reads in the samples, small differences in sequencing library preparation may introduce some additional biases: i) insert sizes may vary between samples, with longer insert sizes leading to a higher power to identify TEs, ii) coverage heterogeneity between samples may be different (e.g. due to different DNA polymerases) and iii) genome sizes may differ between samples (e.g. due to different TE content), where larger genomes will result in lower coverage and thus fewer detected TEs. We address these problems by developing a new data format, the physical pileup track. Analogous to the pileup track, which summarizes for every genomic site the base calls, the physical pileup summarizes for every genomic site the structural states (e.g. TE insert presence or absence). Based on the physical pileup our new software tool PoPoolationTE2 homogenizes the physical coverage across samples and thus also the power to identify TEs.

## PoPoolationTE2

PoPoolationTE2 is a fast and user-friendly tool for analysing TE insertions in one or more samples, where samples could be tissues, pooled populations or sequenced individuals. PoPoolationTE2 does not rely on a set of known TEs in the reference genome, thus both novel and already known TE insertions can be characterized. Compared to its predecessor PoPoolationTE (Kofler et al., 2012), PoPoolationTE2 is designed to compare TE abundance among multiple samples. Additionally, PoPoolationTE2 is substantially faster than its predecessor, as it is implemented in Java and uses bam files as input. It requires paired end data for at least one sample, a reference genome and either a set of TE sequences or a TE annotation. While PoPoolationTE2 accounts for heterogeneity in sequence coverage, the number of chromosomes contributing to the pools should be similar among samples or much larger than the coverage in all samples (minimizing multiple sampling from one individual at a given genomic position).

PoPoolationTE2 requires reads to be mapped to a modified genome, consisting of a reference genome with masked TE sequences and a set of TE sequences. Masking of TEs may be done based on a TE annotation, RepeatMasker (Smit et al., 1996–2010) or, as RepatMasker sometimes misses TE insertions (Rahman et al., 2015), iterative mapping of reads derived from TE sequences (see Manual). When reads are mapped to such a modified genome, TE insertions will result in groups of discordantly mapped paired ends, where one read maps to the reference chromosome and the other to a TE sequence (signatures of TE insertions; fig. 1A). Based on this information a physical pileup track can be created for all samples [fig. 1B; for the difference between base and physical coverage see also Meyerson et al. (2010)]. For every site in the genome, the physical pileup summarizes the structural status inferred from paired end reads. Properly mapped pairs support the absence of a TE while TE insertions are detected by pairs with one read mapping to the reference genome and the other one to a TE sequence. Paired end reads mapping to unrelated genomic positions (e.g. different chromosomes) support structural rearrangements, a feature that can be used for quality filtering. Because the distance between discordantly mapped reads is not know we use the median of the distance between proper pairs instead (inferred for each sample separately). Note that the power to identify TEs scales with the number of mapped reads as well as the distance between the reads, a property that is also captured by the physical coverage track (fig. 1C). To homogenize the power to identify TEs the physical coverage is sampled to equal levels within and between samples. Signatures of TE insertions are then identified as peaks in the physical coverage track and the population frequency of TEs is estimated for the signatures (fig. 1D). Matching pairs of signatures are joined, resulting in a final list of TE insertions. For every TE, PoPoolationTE2 reports the position, the family, the strand and the population frequency in all samples.

**Figure 1:**
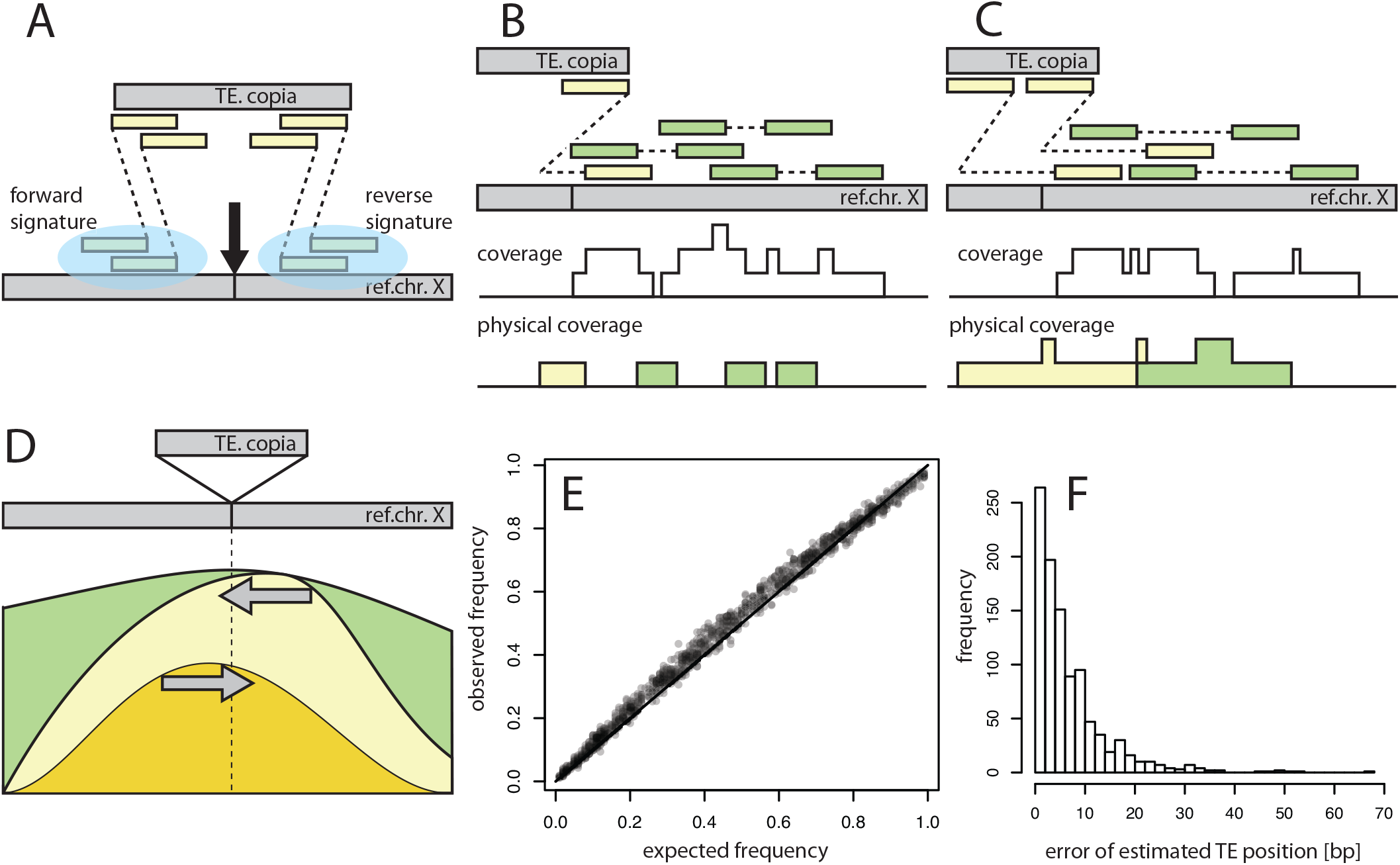
Overview of PoPoolationTE2 A) TE insertions (black arrow) result in paired ends (yellow), with one read mapping to a reference chromosome (X) and the other one to a TE (copia). One group of such discordantly mapped reads is located to the left of the insertion (forward signature) and one to the right (reverse signature). B) PoPoolationTE2 converts the information derived from the position and distance between mapped paired ends into a physical coverage track (bottom). Different types of physical coverage can be derived, for example paired ends mapped as proper pairs (green) or discordant paired ends (yellow) C). Increasing the inner distance between paired ends results in more reads supporting a TE insertion and thus a higher power. In contrast to base coverage which scales only with the number of mapped reads, the physical coverage scales with the number of mapped reads and the inner distance between the reads. D) Signatures of TE insertions (gray arrows) are identified as peaks in the physical pileup track and the population frequency of TEs is computed as ratio between coverage supporting the presence versus absence of a TE. E) Accuracy of the population frequency estimates for 1000 TEs in a simulated pooled population. PoPoolationTE2 has a slight upward bias for intermediate frequency TEs and a slight downward bias for high frequency TEs. F) Accuracy of insertion position estimates for 1000 TEs in a simulated pooled population.

## Performance

We first evaluated the performance of PoPoolationTE2 with simulated data assuming a coverage sufficiently high to detect all TE insertions (supplementary results 1.1). PoPoolationTE2 yields highly reproducible results with different alignment algorithms (bwa mem, bwa bwasw, bowtie2 local), sequencing error rates, read lengths and insert size distributions (supplementary results 1.1). Next, we evaluated the performance for conditions that reflect the properties of typical Pool-Seq studies and found again that PoPoolationTE2 yields reliable results (supplementary results 1.2). The position of TEs is accurately estimated (fig. 1F), but the population frequency of segregating insertions is slightly overestimated and the frequency of fixed insertions is slightly underestimated (fig. 1E).

The crucial question however is, whether PoPoolationTE2 permits an unbiased comparison of TE abundance among samples. To test this we simulated three populations with variable numbers of low frequency (*f* = 0.01) insertions (*A* = 1000, *B* = 750, *C* = 500). For each of these three populations we generated *in silico* different numbers of paired end reads which varied in insert sizes (table 1). With typical Pool-Seq studies only sampling a subset of the chromosomes in the sample (Schlötterer et al., 2014), it is not possible to identify all TE insertions. Nevertheless for an unbiased comparison of different samples it is sufficient to determine the relative TE abundance. Our example in table 1 shows how the analysis of the complete data set may lead to misleading results: in population A fewer TE insertions (minimum count 2) are detected than in population B, despite the opposite being true. Subsampling reads in all samples to equal numbers reduces the problem, but still causes misleading results, with population B again having more insertions (table 1).

**Table 1:** Evaluating different strategies to compare TE abundance in Pool-Seq samples. We simulated three populations with different numbers of low frequency insertions (*f* = 0.01) and paired ends with varying inner distances (ID). An unbiased comparison should result in a stable ratio between observed and simulated TEs in the three populations (i.e. a low *σ_obs/sim_*). The best results were obtained when the physical coverage (phy. cov.) was sampled to equal levels in all three populations. Results are shown for two different minimum count thresholds (mc). The average coverage (*μ_c_*) and the average physical coverage (*μ_pc_*) was directly estimated from the data. * coverage after sampling

| sampling strategy<br>population | naive |  |  | equal reads |  |  | equal phy. cov. |  |  |
| --- | --- | --- | --- | --- | --- | --- | --- | --- | --- |
|  | A | B | C | A | B | C | A | B | C |
| simulated TEs | 1000 | 750 | 500 | 1000 | 750 | 500 | 1000 | 750 | 500 |
| ID | 100 | 150 | 200 | 100 | 150 | 200 | 100 | 150 | 200 |
| reads [million] | 1.045 | 1.379 | 2.045 | 1.045 | 1.045 | 1.045 | 1.045 | 1.379 | 2.045 |
| $\mu_c$ | 199.91 | 266.66 | 399.97 | 199.91 | 202.00 | 204.06 | 199.91 | 266.66 | 399.97 |
| $\mu_{pc}$ | 99.11 | 198.78 | 398.23 | 99.11 | 150.68 | 203.38 | 100.00* | 100.00* | 100.00* |
| observed TEs (mc2) | 395 | 675 | 495 | 397 | 519 | 263 | 150 | 66 | 24 |
| observed/simulated | 0.395 | 0.900 | 0.990 | 0.397 | 0.692 | 0.526 | 0.150 | 0.088 | 0.048 |
| $\sigma_{obs/sim}$ | | 0.321 | | | 0.148 | | | 0.051 | |
| observed TEs (mc1) | 784 | 745 | 496 | 783 | 577 | 264 | 467 | 368 | 253 |
| observed/simulated | 0.784 | 0.993 | 0.992 | 0.783 | 0.769 | 0.528 | 0.467 | 0.491 | 0.506 |
| $\sigma_{obs/sim}$ | | 0.120 | | | 0.143 | | | 0.020 | |

Subsampling the physical coverage to equal levels in all populations consistently resulted in the least biased comparison of TE abundance between populations, irrespective whether paired ends have equal (supplementary table 8) or unequal inner distances (table 1).

Some applications, such as measuring TE activity in mutation accumulation lines, may depend on a reliable identification of sample specific insertions.

This could be challenging as a putative absence of a TE insertion in one sample may in fact be an artefact of coverage heterogeneity. We show that coverage heterogeneity among samples may result in a substantial fraction of false sample specific insertions (supplementary table 9). Restricting the analysis to regions with sufficient physical coverage in all samples dramatically reduces the number of false positives (supplementary table 9).

We conclude that PoPoolationTE2 is a fast and user-friendly tool for an unbiased comparison of TE abundance between samples, thus enabling to study TE dynamics in a broad range of applications.

## Availability

PoPoolationTE2 is implemented in Java and freely available at https://sourceforge.net/projects/popoolation-te2/; For a detailed manual and a walkthrough using a small sample data set see https://sourceforge.net/p/popoolation-te2/wiki/Home/.

## Author’s contributions

RK and CS conceived the study. RK and DGS wrote the software. RK conducted the analysis. RK and CS wrote the paper

## Acknowledgements

We thank all members of the Institute of Population Genetics for feedback and support. This work was supported by the European Research Council Grant ”Archadapt” and the DFG (Adaptomics, SPP 1529).

